# Post-Transcriptional Bone Morphogenetic Protein 2 (BMP2) Gene Regulation in Aorta

**DOI:** 10.1101/735852

**Authors:** Tapan A. Shah, Ying Tang, Edward J. Yurkow, Melissa B. Rogers

**Affiliations:** Rutgers - New Jersey Medical School, Microbiology, Biochemistry, & Molecular Genetics, Newark, NJ; Rutgers University Molecular Imaging Center (RUMIC), Rutgers University, Piscataway, NJ

**Keywords:** BMP, Klotho, gene regulation, signaling, cardiovascular, calcification, atherosclerosis, renal physiology, sex differences

## Abstract

Deletion of an “ultra-conserved sequence” (UCS) within the *Bone Morphogenetic Protein (Bmp)2* mRNA previously revealed that the sequence represses *Bmp2* reporter gene expression in vascular cells. The objective was to determine the impact of the endogenous UCS on *Bmp2* mRNA levels, BMP signaling, and calcification in the healthy control aorta and in the calcified aorta of mice with renal disease. We compared the phenotypes of mice bearing a wild type *Bmp2* allele or the UCS deletion allele in mice with normal kidney function or in *Klotho* mutant mice with reduced kidney function. BMP signaling and calcium levels were normally higher in control females relative to males. UCS deletion induced aortic *Bmp2* mRNA and BMP signaling in control males, but not in females. UCS deletion significantly increased BMP signaling in both male and female *Klotho* homozygotes. Inheritance of the *Bmp2* UCS deletion and *Klotho* alleles was skewed from Mendelian expectations suggesting that these alleles influence interacting pathways. Analyses of body and heart weight supported these interactions. The *Bmp2* UCS represses BMP signaling in control males and in mice of both sexes with abnormal mineralization associated with kidney disease. Disease and sex-specific differences in *Bmp2* gene control may influence the onset and progression of cardiovascular diseases.

## Introduction

The pro-osteogenic bone morphogenetic protein 2 (BMP2) is a potent pro-calcific signal (3–10). Moreover, BMP2 and its downstream effectors, *e.g.*, phosphorylated SMAD1/5/9(8), are implicated strongly in pathological calcification (5,6,9–13). Various parallels exist between osteogenesis and cardiovascular calcification *via* the BMP2 link. BMP2 can induce osteogenic factors in human aortic valve interstitial cells (13) and aortic smooth muscle cells (14). BMP2 induces ossification in diseased aortic valves (9,11,15) and in atherosclerotic plaques (6) and induces calcification *in vitro* (10,13,16–18). Despite an explicit role of Bmp2 in cardiovascular calcification, the mechanisms regulating the patterns of BMP2 and its downstream effectors in the healthy and diseased aorta are incompletely understood. Significant and unanswered questions are: What restrains *Bmp2* expression in healthy cardiovascular tissues? Why is *Bmp2* induced in physiologies such as aging and reduced renal function that promote pathological calcification? Elucidating the mechanisms that control BMP2 synthesis leading to pathological calcification may reveal new therapeutic strategies.

Several *cis* and *trans*-acting factors can regulate *Bmp2* gene expression either transcriptionally or post-transcriptionally [reviewed in (19) and (20)]. The *Bmp2* gene may be “active”; *i.e.*, transcribed, but a post-transcriptional block may prevent BMP2 synthesis in specific cell types. Our studies showed that a unique ultra-conserved sequence (UCS) in the 3’ untranslated region (UTR) of the transcript mediates this repression (21–23). The UCS repressed reporter genes in mesenchymal and other types of non-transformed cells *in vitro* (24–27) as well as in the coronary vasculature, valves, and the aorta *in vivo* (25,27). The fact that these tissues are prone to calcification in patients with CAVD and atherosclerosis risk factors, suggests the hypothesis that UCS-mediated repression may protect against pathological levels of BMP2 synthesis leading to calcification.

Our previous findings in embryos demonstrated that the UCS limits *Bmp2* mRNA abundance and BMP signaling and that disturbing *Bmp2* 3’UTR-mediated events negatively impacted embryonic development. Here we describe the impact of an allele lacking the UCS (*Bmp2*^Δ*UCS*^) on *Bmp2* expression and BMP signaling in the adult aorta of healthy control mice and in mice with a mutation that causes renal failure and premature aging.

BMP2 levels are normally low in the healthy vasculature but are induced in pathologically calcified tissues. Mutation of the *Klotho* gene provides an experimental model in which we could compare the mechanisms that repress or activate *Bmp2*. KLOTHO is a protein largely synthesized in the kidney that controls mineral metabolism. KLOTHO deficiency promotes hyperphosphatemia that leads to rapid and dramatic calcification of the aorta and aortic valve by 6-7 weeks of age (28,29). In contrast, calcification in other models, *e.g.*, hyperlipidemia, is quite slow (30). Furthermore, BMP2 protein was observed in the calcified aortic valves of *Klotho* null mice and BMP signaling was shown to be required for aortic valve calcification (29). In this study, we describe how the UCS affects *Bmp2* RNA abundance, BMP signaling, and vascular calcification in the aorta and the effect on the overall fitness of control mice and *Klotho* mutant mice of both sexes.

## Materials and Methods

### Mouse strains

The background of all mice bearing the *Bmp2* and *Klotho* mutations was a mixture of strains 129, C57Bl/6J, and C3H/J. The UCS deletion allele (*Bmp2*^*ΔUCS*^) was described in Shah et al. (31). Mice bearing the *Klotho* mutation (28) were a gracious gift from Dr. Makoto Kuro-o (Jichi Medical University) by way of Dr. Sylvia Christakos (Rutgers New Jersey Medical School). C57Bl/6 female mice (47 days, 6 months, 12 months, 18 months and 22 or 23 months old) and male mice (47 days, 6 months, 12 months, 18 months and 21 months old) were obtained from the National Institute of Aging (NIA) aged rodent colonies (Bethesda, MD).

### Mice Handling

Animals were handled in accordance with the Guidelines for Care and Use of Experimental Animals and approved by the NJ Medical School Institutional Animal Care and Use Committee (IACUC protocol #15069). Control and *Klotho* homozygote mice were fed regular chow and euthanized at 47 ± 6 days of age. After weaning, *Klotho* homozygotes received softened chow on the floor of the cage. Aortas were obtained from 10-week-old C57Bl/6 females that were ovariectomized for another study (Khariv and Elkabes in preparation, IACUC protocol #15038). Briefly, mice were anesthetized and both ovaries surgically removed. Sham mice underwent surgery without removal of ovaries. Three weeks post-surgery, mice were euthanized and necropsied as described below.

On the day of necropsy, mice were killed with an inhalation overdose of isoflurane. Immediately thereafter, the heart was perfused via the left ventricle with phosphate buffered saline (PBS, pH 7.3), to remove excess blood. The heart and aorta down to the aortic abdominal bifurcation into the left and right common iliac arteries were removed intact. After cutting the aorta at the surface of the heart, both tissues were rinsed in PBS, blot dried, and weighed. The heart was fixed in neutral buffered formalin. The aorta was snap-frozen in liquid nitrogen and stored at −80°C. For biochemical assays, frozen tissues were ground in liquid nitrogen using a mortar and pestle. The frozen powder was split to be used for different assays. To facilitate the handling of small tissues such as the diseased *Klotho* aortas, glass beads (Millipore-SIGMA, St. Louis, MO, # G1277) were added during the grinding. Glass beads did not affect the biochemical assays (Fig. 1A, B).

**Figure 1.**
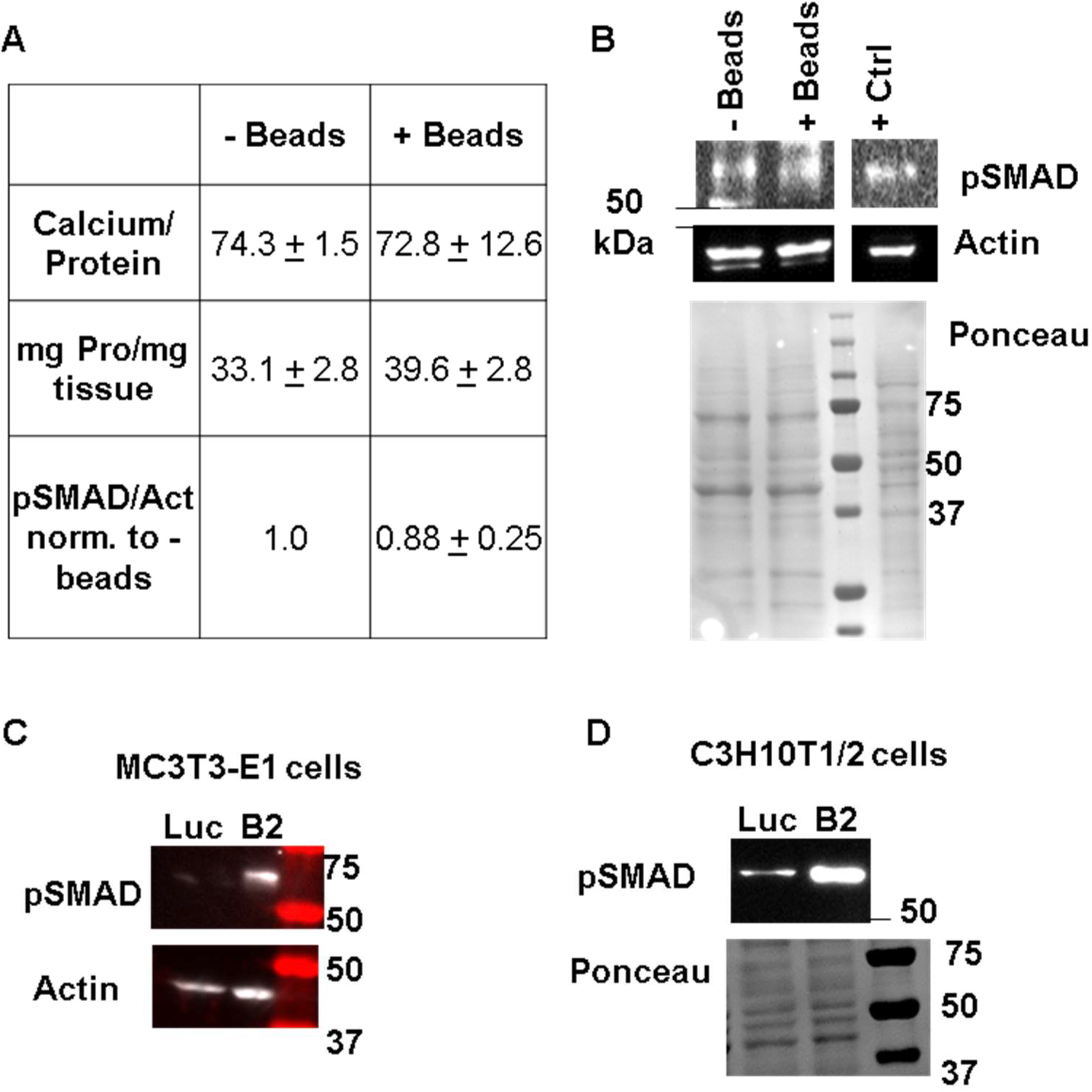
Experimental Controls. **A, B.** Grinding tissue with glass beads does not significantly affect calcium or protein yield. Whole aorta was ground in liquid nitrogen either with or without glass beads. Frozen ground powder tissue was split two ways for protein and calcium assays. **A.** Effect of glass beads on calcium, protein and pSMAD 1/5/9(8) levels. Duplicate measurements are presented with range. Experiments were repeated twice with similar results. **B.** Representative blots showing pSMAD 1/5/9(8), actin levels and the Ponceau S stained membrane after transfer. The positive control lane (+ Ctrl) was loaded with lysate from MC3T3-E1 cells transfected with a *Bmp2* expression plasmid. **C, D.** Validation of the phospho-SMAD 1/5/9(8) antibody. MC3T3-E1 **(C)** and C3H10T1/2 **(D)** cells were transfected with an expression plasmid encoding *Bmp2* (B2) or luciferase (Luc) (25). Cells were then lysed in RIPA buffer and subjected to western blotting as described in the experimental procedures section. These representative blots show that pSMAD1/5/9(8) levels were induced in cells transfected with the Bmp2 expressing plasmid relative to the luciferase plasmid. Actin levels and a Ponceau S stained membrane are shown as loading controls.

### Genotyping

Genomic DNA was isolated and *Bmp2* genotypes were determined by semi-quantitative PCR as described in Shah *et al* (31). *Klotho* genotypes were determined by semi-quantitative PCR using LA-Tag DNA polymerase and TaKaRa buffer with Mg^+2^ (TaKaRa Shuzo, Tokyo, Japan) as follows: initial denaturation at 97°C for 2 min; followed by 32 cycles of denaturation at 94°C for 30 secs; annealing at 55°C for 30 secs; and extension at 72°C for 1 min 30 secs; ending with a final extension at 72°C for 10 min. A common primer (TGGAGATTGGAAGTGGACG, 0.2 μM final concentration) and a wild type specific primer (TTAAGGACTCCTGCATCTGC, 0.05 μM final concentration) amplified a 458 bp fragment from the wild type *Klotho* allele. The common primer and a mutation specific primer (CAAGGACCAGTTCATCATCG, 0.2 μM final concentration) amplified a 920 bp fragment from the *Klotho* mutant allele (32). Some *Bmp2* and *Klotho* genotyping was performed by Transnetyx, Inc, Cordova, TN.

### RNA Isolation and Reverse Transcription and Quantitative Real Time PCR (RT qPCR)

Total RNA was isolated using the miRNeasy kit (Qiagen Inc., Germantown, MD, # 217004). RNA quantity and quality (A260/280) were determined using a NanoDrop spectrophotometer (NanoDrop Technologies, Wilmington, DE). cDNA was synthesized using the QuantiTect® Reverse Transcription kit (Qiagen Inc., Germantown, MD, # 205313). Quantitative PCR was performed using the QuantiTect® SYBR® Green PCR kit (Qiagen Inc., Germantown, MD, #204145) and a CFX96 Touch™ Real-Time PCR Detection System (Bio-Rad Laboratories, Hercules, CA, #1855196). Relative *Bmp2* mRNA expression was calculated using CFX96 Manager software (Bio-Rad Laboratories, Hercules, CA, # 1845000) with actin as the reference gene. Intron-spanning primers were used to eliminate amplicons generated from any contaminating genomic DNA. The primer sequences used were *Bmp2* - Forward (TAGATCTGTACCGCAGGCA) and Reverse (GTTCCTCCACGGCTTCTTC) and *Actin* - Forward (CGCCACCAGTTCGCCATGGA) and reverse (TACAGCCCGGGGAGCATCGT).

### Western blots

Frozen ground tissue was solubilized in RIPA buffer, sonicated, and subjected to western blot analyses as described in Shah et al (31). BMP signaling was measured using a monoclonal phospho-SMAD 1/5/9(8) antibody (Cell Signaling Technology, Danvers, MA, #13820) at a dilution of 1:1000. The pSMAD antibody was authenticated as described in (31) and Fig. 1C, D. Polyclonal total SMAD 1/5/9(8) (Santa Cruz Biotechnology, Inc., Santa Cruz, CA, #sc-6031-R) and a polyclonal actin antibody (Santa Cruz Biotechnology, Inc., Santa Cruz, CA, #sc-1615-R) were subsequently used at a dilution of 1:1000. In all cases, the secondary antibody was Goat Anti-Rabbit HRP (Abcam, Cambridge, MA, # ab97080) at a dilution of 1: 20,000. Antibody-bound proteins were detected using SuperSignal™ West Femto Maximum Sensitivity Substrate (ThermoFisher Scientific, Waltham, MA, # 34096) and imaged using a FluoroChem M (Protein Simple, San Jose, California).

### Calcium assays

Ground tissue was solubilized and lysed by sonication on ice in PBS, pH 7.3 containing 0.16 mg/mL heparin. The Cayman Chemical Calcium Assay kit was used to measure calcium levels (Ann-Arbor, MI, #701220). Calcium levels were normalized to protein levels measured using the Bradford assay (Bio-Rad Laboratories, Hercules, CA, # 5000006).

### Spatial mapping of calcified structures

The patterns of mineralization in *Klotho* heterozygote *vs. Klotho* mutant mice were determined using microcomputerized tomography (microCT) at the Rutgers Molecular Imaging Center (http://imaging.rutgers.edu/). Mice were scanned using the Albira® PET/CT (Carestream, Rochester, NY) at standard voltage and current settings (45kV and 400μA) with a minimal voxel size of <35 μm. Voxel intensities in the reconstructed images were evaluated and segmented with VivoQuant image analysis software (version 1.23, inviCRO LLC, Boston).

### Statistical Analysis

The statistical significance was determined using student’s *t* test, Chi-square test, linear regression and two-way ANOVA analysis. A *p* value of less 0.05 was considered statistically significant.

## Results

### Adult mice lacking the *Bmp2* UCS

We previously demonstrated that the *Bmp2* UCS represses reporter gene expression *in vitro* in mesenchymal cells (20,24,25) and *in vivo* in the aorta and coronary vasculature (25). These tissues are prone to pathological calcification. Furthermore, we showed that the *Bmp2* UCS represses *Bmp2* RNA abundance and BMP signaling in mid-gestation embryos (31). Based on these findings, we hypothesized that the *Bmp2* UCS represses *Bmp2* RNA levels, BMP signaling and calcium levels in the aorta. Although mice completely lacking the UCS were underrepresented (31), some mice did survive to adulthood. Consequently, we tested the impact of the UCS in the aorta of these adults. In all cases, both sexes were assayed, because sex hormones can influence the expression of various members of the BMP signaling pathway (33). We also tested UCS function in pathologically calcified aorta from mice with renal failure and premature aging due to KLOTHO deficiency (28). To obtain aorta from control or diseased mice with three *Bmp2* genotypes (wild type *Bmp2*^+/+^, heterozygous *Bmp2*^+/Δ*UCS*^, or homozygous *Bmp2*^Δ*UCS*/Δ*UCS*^ for the UCS deletion), we mated parents that were heterozygous for the *Bmp2* allele lacking the UCS (*Bmp2*^+/*ΔUCS*^) and for the *Klotho* mutant allele (*Kl*^*kl*/+^). The expected and actual fractions of each genotype are shown in Fig. 2A. In mice with adequate KLOTHO levels (*Kl*^+/+^ or *Kl*^*kl*/+^), no statistically significant skewing from the Mendelian inheritance of the *Bmp2* mutant allele was observed (Fig. 2B).

**Figure 2.**
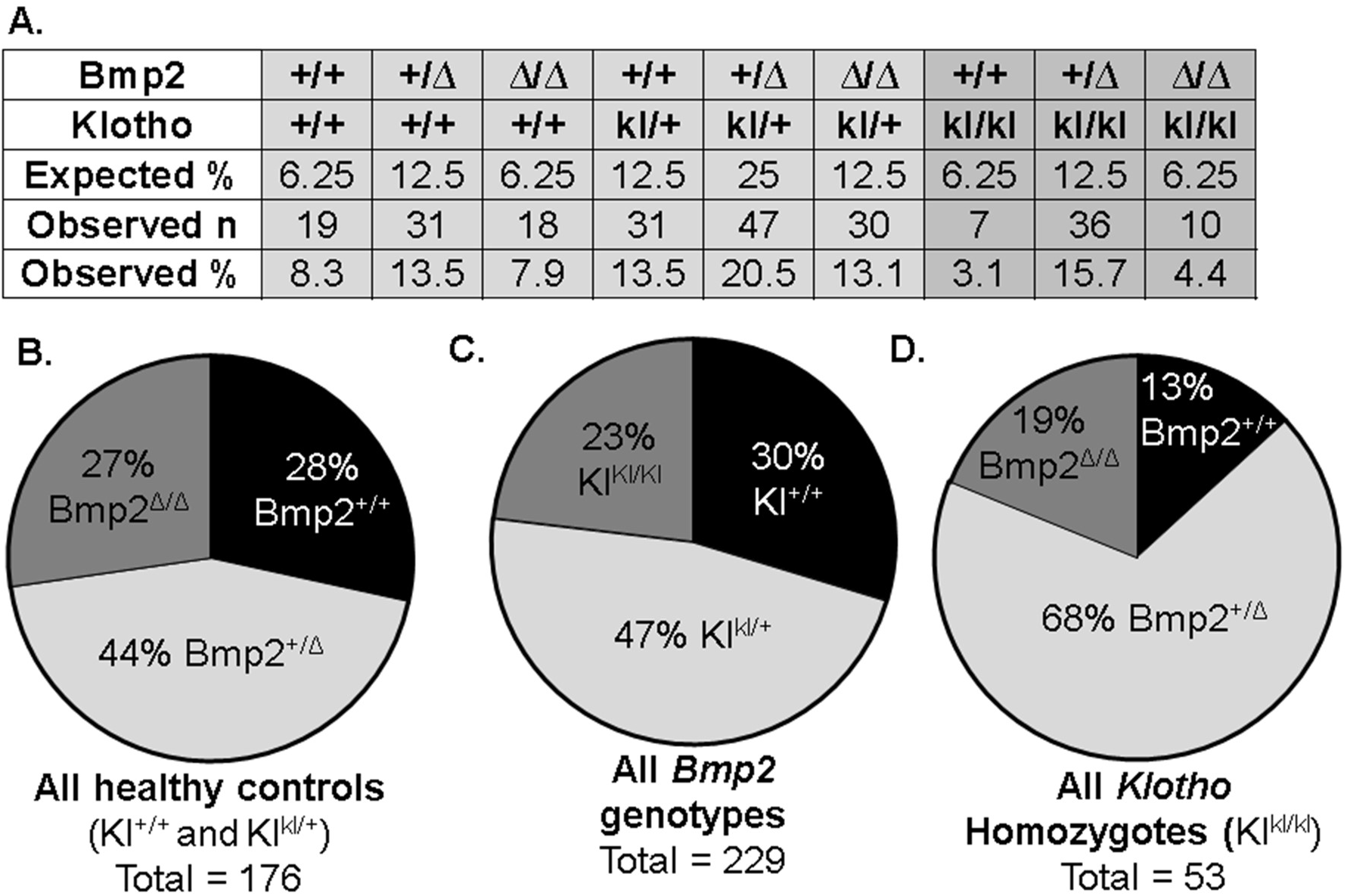
Apparent interaction between *Bmp2* and *Klotho* mutant alleles. Mice heterozygous for the *Bmp2* allele lacking the UCS (*Bmp2*^+/*ΔUCS*^) and for the *Klotho* mutant allele (*Kl*^*kl*/+^) were bred. The resulting pups were genotyped prior to weaning at approximately 4 weeks of age. *Bmp2* genotypes: wild type (+/+), heterozygous (+/Δ), or homozygous UCS deletion (ΔΔ) and *Klotho* genotypes: wild type (+/+), heterozygous (kl/+), or homozygous *Klotho* mutation (kl/kl). **A.** Expected and observed genotypes for the breeding scheme described above. Occasionally, pups were found dead between birth and weaning, but the frequency did not differ from that typically observed, nor was any particular genotype obviously over-represented. **B-D** illustrate relative fractions for each genotype.

The *Klotho* mutant allele is recessive. The first visible phenotype of homozygous *Klotho* mutant mice (*Kl*^*kl*/*kl*^) is runting relative to control littermates at the time of weaning. Although significant prenatal or perinatal lethality has not been reported, we observed underrepresentation of *Klotho* homozygotes, although the difference from Mendelian expectations was not quite significant (Fig. 2C). However, segregation of the *Bmp2* genotypes was significantly skewed in the *Klotho* homozygotes (Fig. 2D) with a significant overrepresentation of heterozygotes (*Bmp2*^+/Δ*UCS*^) bearing one wild type and one UCS deletion allele (*Bmp2*^+/Δ*UCS*^, Chi-squared equals 7.15 with 2 degrees of freedom, two-tailed *p*-value < 0.03). These results suggest that changes in BMP2 levels associated with this regulatory mutation alter the fitness of the mice with KLOTHO deficiency. Indeed, as will be discussed below, *Bmp2* genotype was associated with significant differences in weight specifically in the diseased *Klotho* homozygotes.

To confirm that the *Klotho* mutant allele is fully recessive, we compared wild type (*Kl*^+/+^) mice and heterozygotes that inherited one wild type and one mutated *Klotho* allele (*Kl*^*kl*/+^). We found that the following parameters: *Bmp2* RNA levels, BMP signaling, calcium levels, and organ and body weights did not differ between *Klotho* heterozygotes and mice with two wild type *Klotho* alleles (Table 1). Therefore, the results from *Klotho* “control” mice presented below include data from both wild type and *Klotho* heterozygotes. The influence of *Bmp2* genotype will be discussed first in control mice and then in the diseased homozygotes.

**Table 1.** *Klotho* wild type and *Klotho* heterozygous mice are equivalent controls. Two-way ANOVA analysis was performed to confirm that there was no significant difference between *Klotho* wildtype (*Kl*^+/+^) and *Klotho* mutant heterozygotes (*Kl*^*kl*/+^) mice. Parameters compared included *Bmp2* RNA, pSMAD normalized to total SMAD, aortic calcium levels, body weight, heart weight, heart weight normalized to body weight and age at euthanasia. The average (Avg.), standard deviation (SD), and number of mice assayed (n) is shown.

|  | MALES |  |  |  |  |  |  | FEMALES |  |  |  |  |  |
| --- | --- | --- | --- | --- | --- | --- | --- | --- | --- | --- | --- | --- | --- |
| Bmp2 genotype | Wt |  |  |  |  |  |  | Wt |  |  |  |  |  |
| Klotho genotype | Wt |  |  | K+ |  |  |  | Wt |  |  | K+ |  |  |
| Two-way Anova<br>(p val) | 0.29 |  |  |  |  |  |  | 0.21 |  |  |  |  |  |
| Parameters | Avg. | SD | n | Avg. | SD | n |  | Avg. | SD | n | Avg. | SD | n |
| Bmp2 RNA | 0.72 | 0.24 | 7 | 0.78 | 0.46 | 16 |  | 0.57 | 0.09 | 5 | 0.89 | 0.31 | 6 |
| pSMAD/Total SMAD | 0.61 | 0.30 | 4 | 0.58 | 0.05 | 2 |  | 1.16 | 0.50 | 3 | 1.15 | 0.79 | 4 |
| Calcium/Protein | 42.1 | 21.8 | 7 | 51.2 | 21.2 | 11 |  | 64.8 | 23.4 | 5 | 53.2 | 14.5 | 3 |
| Body Weight (g) | 22.3 | 2.8 | 13 | 22.3 | 1.4 | 18 |  | 18.2 | 1.6 | 8 | 18.8 | 2.3 | 10 |
| Heart Weight (g) | 0.097 | 0.027 | 13 | 0.111 | 0.021 | 16 |  | 0.082 | 0.014 | 8 | 0.093 | 0.013 | 10 |
| Heart / Body Weight | 0.0044 | 0.0012 | 13 | 0.0049 | 0.0010 | 15 |  | 0.0045 | 0.0004 | 8 | 0.0049 | 0.0006 | 10 |
| Age (d) at euthanasia | 47.8 | 3.7 | 13 | 48.9 | 1.8 | 19 |  | 49.4 | 3.1 | 8 | 48.5 | 3.5 | 10 |
|  | MALES |  |  |  |  |  |  | FEMALES |  |  |  |  |  |
| Bmp2 genotype | +/ΔUCS | | | | | | | +/ΔUCS | | | | | |
| Klotho genotype | Wt |  |  | K+ |  |  |  | Wt |  |  | K+ |  |  |
| Two-way Anova<br>(p val) | 0.96 |  |  |  |  |  |  | 0.81 |  |  |  |  |  |
| Parameters | Avg. | SD | n | Avg. | SD | n |  | Avg. | SD | n | Avg. | SD | n |
| Bmp2 RNA | 0.98 | 0.48 | 10 | 1.02 | 0.45 | 11 |  | 0.66 | 0.36 | 3 | 1.17 | 0.34 | 6 |
| pSMAD/Total SMAD | 1.00 | 0.30 | 5 | 1.00 | 0.06 | 3 |  | 1.00 | 0.61 | 5 | 1.00 | 0.33 | 3 |
| Calcium/Protein | 50.1 | 22.1 | 10 | 50.1 | 25.2 | 16 |  | 63.6 | 30.0 | 7 | 66.6 | 18.5 | 11 |
| Body Weight (g) | 22.4 | 3.0 | 16 | 21.9 | 3.3 | 26 |  | 19.5 | 3.2 | 15 | 19.3 | 2.2 | 22 |
| Heart Weight (g) | 0.11 | 0.02 | 15 | 0.11 | 0.02 | 23 |  | 0.09 | 0.02 | 11 | 0.10 | 0.02 | 13 |
| Heart / Body Weight | 0.0051 | 0.0007 | 15 | 0.0051 | 0.0012 | 22 |  | 0.0045 | 0.0010 | 11 | 0.0052 | 0.0008 | 12 |
| Age (d) at euthanasia | 46.9 | 5.8 | 16 | 46.9 | 6.2 | 27 |  | 49.1 | 2.6 | 15 | 48.5 | 5.9 | 23 |
|  | MALES |  |  |  |  |  |  | FEMALES |  |  |  |  |  |
| Bmp2 genotype | ΔUCS/ΔUCS |  |  |  |  |  |  | ΔUCS/ΔUCS |  |  |  |  |  |
| Klotho genotype | Wt |  |  | K+ |  |  |  | Wt |  |  | K+ |  |  |
| Two-way Anova<br>(p val) | 0.26 |  |  |  |  |  |  | 0.52 |  |  |  |  |  |
| Parameters | Avg. | SD | n | Avg. | SD | n |  | Avg. | SD | n | Avg. | SD | n |
| Bmp2 RNA | 1.71 | 0.16 | 4 | 1.48 | 0.58 | 8 |  | 2.12 | 0.75 | 2 | 0.65 | 0.26 | 8 |
| pSMAD/Total SMAD | 1.20 | 0.17 | 4 | 1.43 | 0.34 | 3 |  | 1.10 | 0.33 | 3 | 0.87 | 0.74 | 3 |
| Calcium/Protein | 36.1 | 11.0 | 8 | 45.6 | 18.3 | 11 |  | 67.9 | 27.7 | 4 | 60.5 | 30.7 | 9 |
| Body Weight (g) | 22.9 | 2.6 | 15 | 22.7 | 3.5 | 23 |  | 18.9 | 2.4 | 12 | 18.2 | 1.6 | 24 |
| Heart Weight (g) | 0.115 | 0.014 | 11 | 0.114 | 0.021 | 15 |  | 0.099 | 0.023 | 8 | 0.098 | 0.019 | 19 |
| Heart / Body Weight | 0.0051 | 0.0008 | 11 | 0.0052 | 0.0006 | 15 |  | 0.0052 | 0.0010 | 8 | 0.0054 | 0.0011 | 18 |
| Age (d) at euthanasia | 47.7 | 4.4 | 15 | 47.3 | 5.5 | 23 |  | 47.5 | 5.6 | 12 | 48.7 | 5.4 | 25 |

### The *Bmp2* UCS influences *Bmp2* RNA and BMP signaling levels in control aorta

We first tested the effect of UCS deletion in aorta from healthy control mice. Using RT qPCR, we measured relative *Bmp2* RNA levels in aorta from mice that were wild type, heterozygous, or homozygous for the *Bmp2* UCS deletion allele. Fig. 3A shows that aortic *Bmp2* RNA abundance increased with the number of UCS deletion alleles in males. Specifically, aortic *Bmp2* RNA abundance was 1.3-fold higher in heterozygotes (*Bmp2*^+/Δ*UCS*^) and 2-fold higher in homozygotes (*Bmp2*^Δ*UCS*/Δ*UCS*^, *p* = 2.6 × 10^−5^) relative to wild type mice (*Bmp2*^+/+^). Interestingly, although the UCS acted as a repressor in males, UCS deletion did not affect *Bmp2* RNA abundance in females (Fig. 3A).

**Figure 3.**
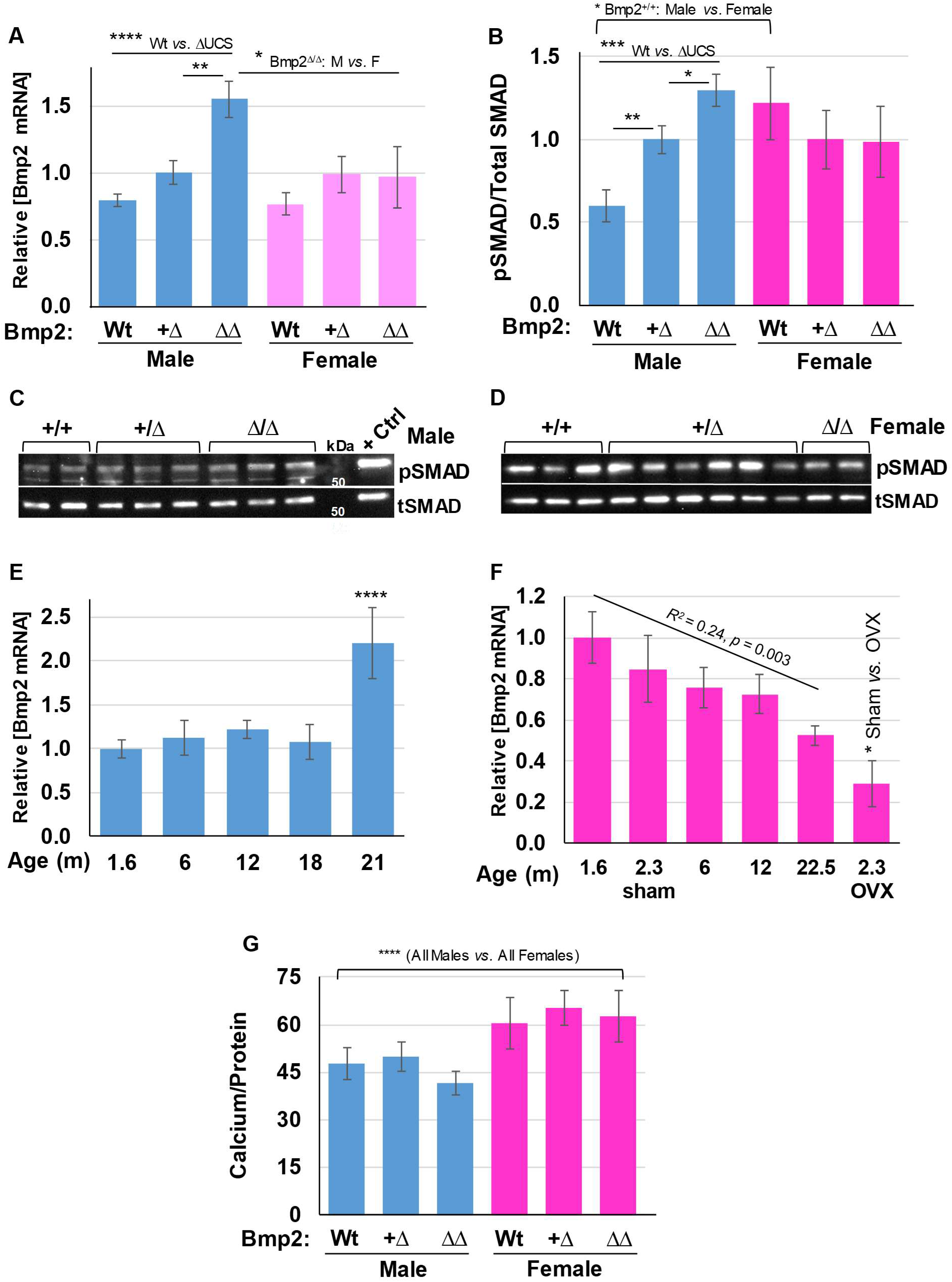
*Bmp2* RNA, BMP signaling, and calcium levels in aorta from control mice with different *Bmp2* genotypes. Aortas were isolated from mice bearing the wild type (Wt), heterozygous (+/Δ), or homozygous UCS deletion (ΔΔ) *Bmp2* genotypes between 40 and 50 days of age. Mice were healthy with normal kidney function (*Kl*^+/+^ and *Kl*^*kl*/+^). Average parameter values are presented with the standard error of the mean (SEM). **A.** *Bmp2* RNA levels normalized to actin RNA levels (males, n = 12 – 21; females, n = 9 – 11). **B.** BMP signaling levels as assessed by phosphorylated SMAD1/5/9(8) (pSMAD) levels normalized to total SMAD 1/5/9(8) (tSMAD) levels (males, n = 6 – 8; females, n = 6 – 8). **C, D.** Representative western blot panels showing pSMAD1/5/9(8) and total SMAD1/5/9(8) levels, males (left) and females (right). The positive control lane (+ Ctrl) was loaded with lysate from MC3T3-E1 cells transfected with a *Bmp2* expression plasmid (Fig. 1). **E, F.** *Bmp2* RNA levels normalized to actin RNA levels in the aorta from male and female mice (ages below each bar, n = 4 - 6) and ovariectomized (OVX) and sham control mice (euthanized at 69 days, 3 weeks after operation, n = 3). **G.** Calcium levels normalized to protein concentration (males, n = 18 – 26; females, n = 6 – 18). All experiments were repeated at least twice with similar results. * *p* < 0.05, ** *p* < 0.01, \*\*\**p* < 0.005, **** *p* < 0.001.

We then measured the impact of UCS deletion on BMP signaling as assessed by the phosphorylation of SMAD1/5/9(8), the canonical BMP signaling intermediates, using an antibody authenticated as described previously ((31), Fig. 1C, D). The ratio of phosphorylated protein pSMAD1/5/9(8) relative to total SMAD1/5/9(8) levels was 1.7-fold higher in aorta from male heterozygotes (*Bmp2*^+/Δ*UCS*^, *p* = 0.008) and 2.2-fold higher in male homozygotes (*Bmp2*^Δ*UCS*/Δ*UCS*^, *p* = 3.5 × 10^−4^) relative to wild type male mice (Fig. 3B, C). These results are consistent with the hypothesis that the *Bmp2* UCS represses *Bmp2* synthesis in the aorta. In contrast to males, deleting the *Bmp2* UCS failed to induce either *Bmp2* RNA or BMP signaling in females (Fig. 3A - C). This result and those described below suggest that sex influences *Bmp2* gene expression.

Estrogen was previously shown to directly induce *Bmp2* transcription in cultured bone marrow mesenchymal stem cells (34). However, the impact of estrogen on *Bmp2* expression has not been tested *in vivo*. To test whether or not this sex steroid impacts *Bmp2* RNA levels in the aorta, we measured *Bmp2* mRNA levels in aged females who do not undergo the pre-ovulatory rise in circulating estrogen levels (35,36). A simple linear regression calculation determined that aortic *Bmp2* RNA levels declined with increasing age (Fig. 3F, R^2^ = 0.35, *p* = 0.003). We also observed reduced *Bmp2* RNA levels in aorta from ovariectomized (OVX) mice relative to mice subjected to a sham operation (Fig. 3F, *p* = 0.05). Interestingly, in males, *Bmp2* mRNA abundance was significantly higher in aorta from 21 months old mice relative to younger male mice (Fig. 3E, *p* = 8.5 × 10^−5^) or similarly aged females (*p* = 0.02). These observations are consistent with both transcriptional and post-transcriptional differences in the mechanisms that regulate BMP2 synthesis in males and females.

We then measured calcium levels normalized to protein concentration in whole aorta lysate from control males and females. We observed that UCS deletion did not elevate calcium levels in aorta from either sex (Fig. 3G). This suggests that regulatory mechanisms in the osteogenic pathway between BMP and calcium deposition may counter the pro-calcific effect of increased BMP signaling. Intriguingly, BMP signaling (Fig. 3B, *p* = 0.04) and calcium levels (Fig. 3G, *p* = 0.0004) in control aorta from females were higher than that in control aorta from males. In summary, sex-dependent differences in the regulatory mechanisms that control aortic calcification include increased BMP signaling and basal calcification levels in females. These increases are accompanied by an apparent lack of UCS-associated repression in females.

### *Klotho* mutant mice: an inducible model of *Bmp2* gene expression

Increased BMP2 mRNA levels and BMP signaling are required for calcification in the aortic valve of mice homozygous for the *Klotho* null allele (29). We tested the role of BMP2 and BMP signaling in the calcified aorta of mice homozygous for the original hypomorphic *Klotho* allele (28). First, we confirmed that *Klotho* mRNA levels in the kidneys of mice homozygous for the *Klotho* mutation were less than 1% of that observed in healthy control mice (Fig. 4A, p = 0.0009). *Bmp2* genotype did not significantly alter *Klotho* RNA abundance (Fig. 4A). BMP signaling and calcium levels in aorta from control mice that were either wild type (*Kl*^+/+^) or heterozygous (*Kl*^*kl*/+^) for the *Klotho* mutation were compared to the calcified aorta from *Klotho* mutant homozygotes (*Kl*^*kl*/*kl*^). BMP signaling was induced nearly 2-fold in aorta from male and female *Klotho* homozygous mice relative to control mice (Fig. 4B, C; *p* = 0.03). Calcium levels in the aorta from *Klotho* mutant homozygotes were significantly elevated by over 2-fold in males (*p* = 1.4 × 10^−6^) and females (*p* = 5.0 × 10^−4^) relative to control aorta (Fig. 4D). PET-CT imaging revealed profound mineralization in the aortic sinus and ascending aorta of *Klotho* mutant (*Kl*^*kl*/*kl*^) mice, but not in a control heterozygous (*Kl*^*kl*/+^) littermate (Fig. 4E, F). Together, these results confirmed that homozygosity for the *Klotho* mutation amplifies BMP signaling and calcification in the aorta.

**Figure 4.**
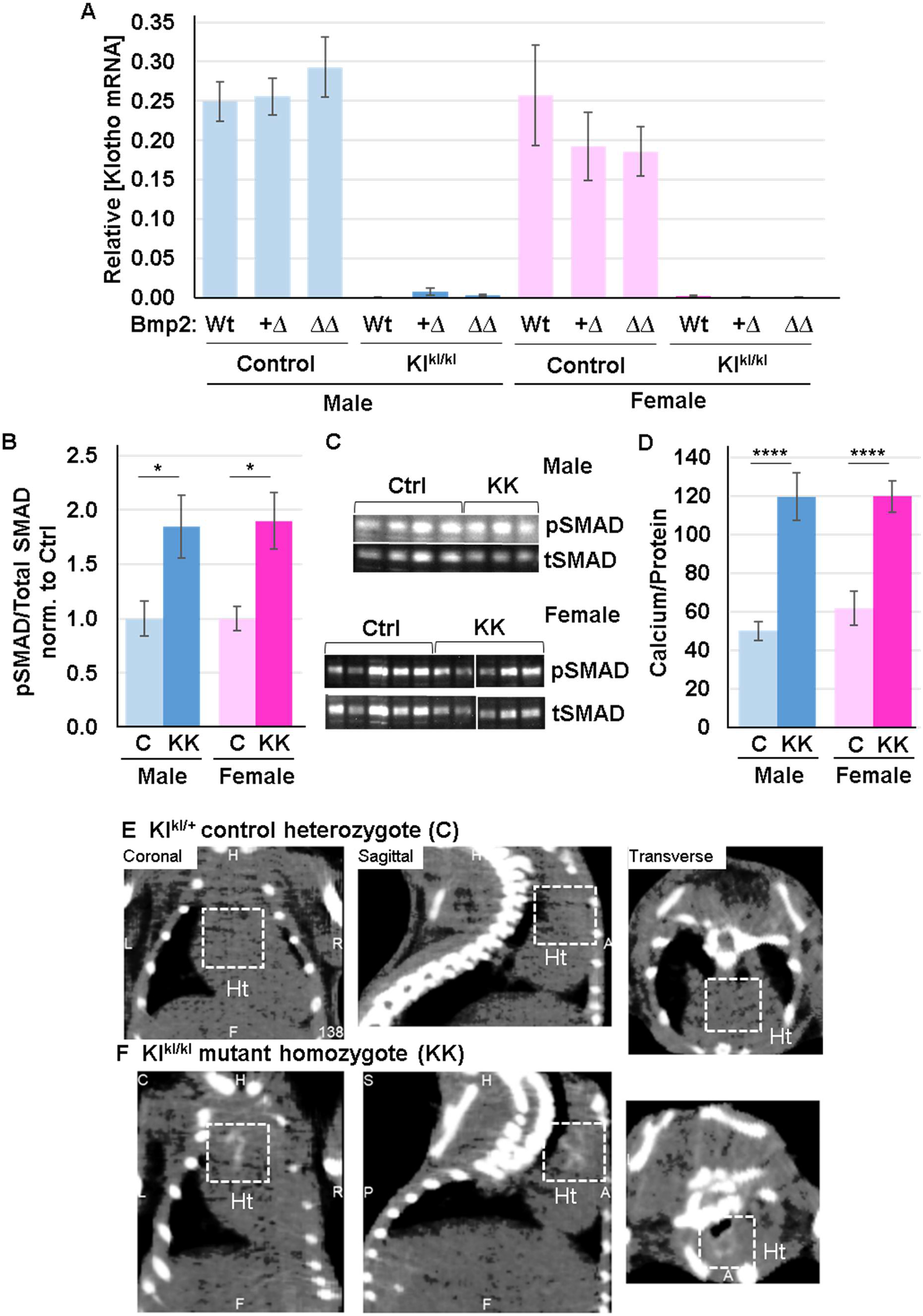
*Klotho* RNA, BMP signaling levels and calcification in aorta from homozygous *Klotho* mutant mice. Aortas were isolated from control (C) mice with normal kidney function (*kl*^+/+^ and *kl*^*kl*/+^) or mice homozygous for the *Klotho* mutation with renal disease (KK) between 40 and 50 days of age. Average parameter values are presented with SEM. **A.** *Klotho* RNA levels normalized to actin RNA levels (males, n = 4 – 18; females, n = 3 – 8). *Bmp2* genotypes are indicated below each bar. **B.** BMP signaling levels as assessed by phosphorylated SMAD1/5/9(8) (pSMAD) levels normalized to total SMAD 1/5/9(8) (tSMAD) (males, n = 3 – 6; females, n = 5 – 7). All mice were wild type for *Bmp2* genotype. **C.** Representative western blot panels showing pSMAD1/5/9(8) and total SMAD1/5/9(8) levels, males (top) and females (bottom). **D.** Average calcium levels normalized to protein concentration (males, n = 9 – 18; females, n = 6 – 7). All experiments were repeated at least twice with similar results. * *p* < 0.05, **** *p* < 0.001. **E, F.** Male littermates were scanned with an Albira PET/CT Imaging System (Carestream, Rochester, NY) set at 45 kV, 400 μA, and <35 μm voxel size. Voxel intensities in the reconstructed images were evaluated and segmented with VivoQuant image analysis software (version 1.23, inviCRO LLC, Boston MA). The dashed white lines mark mineralized areas of the aortic sinus and ascending aorta present in the heart (**Ht**) of the *Klotho* mutant homozygote **(F)**, but not in the heterozygous control *Kl*^*kl*/+^ **(E)**.

### The *Bmp2* UCS represses BMP signaling levels in calcified aorta from *Klotho* mutant mice

We showed earlier that the *Bmp2* UCS repressed mRNA abundance and BMP signaling in non-calcified control aorta from males, but not females (Fig. 3A - C). Consistent with our findings in control aorta from males (Fig. 3B, C), UCS deletion significantly stimulated aortic BMP signaling in male *Klotho* homozygotes (*Kl*^*kl*/*kl*^). BMP signaling was about 2-fold higher in heterozygotes (*Bmp2*^+/Δ*UCS*^, *p* = 0.01) and homozygotes (*Bmp2*^Δ*UCS*/Δ*UCS*^, *p* = 0.02) relative to wild type (*Bmp2*^+/+^) mice (Fig. 5A, B). In contrast to control aorta from females, UCS deletion also stimulated aortic BMP signaling by 1.6-fold in female *Klotho* homozygotes relative to wildtype females (Fig. 5A, B; *p* = 0.02). The observed elevation in BMP signaling did not lead to an obvious stimulation of calcium levels in mice lacking the UCS (Fig. 5C).

**Figure 5.**
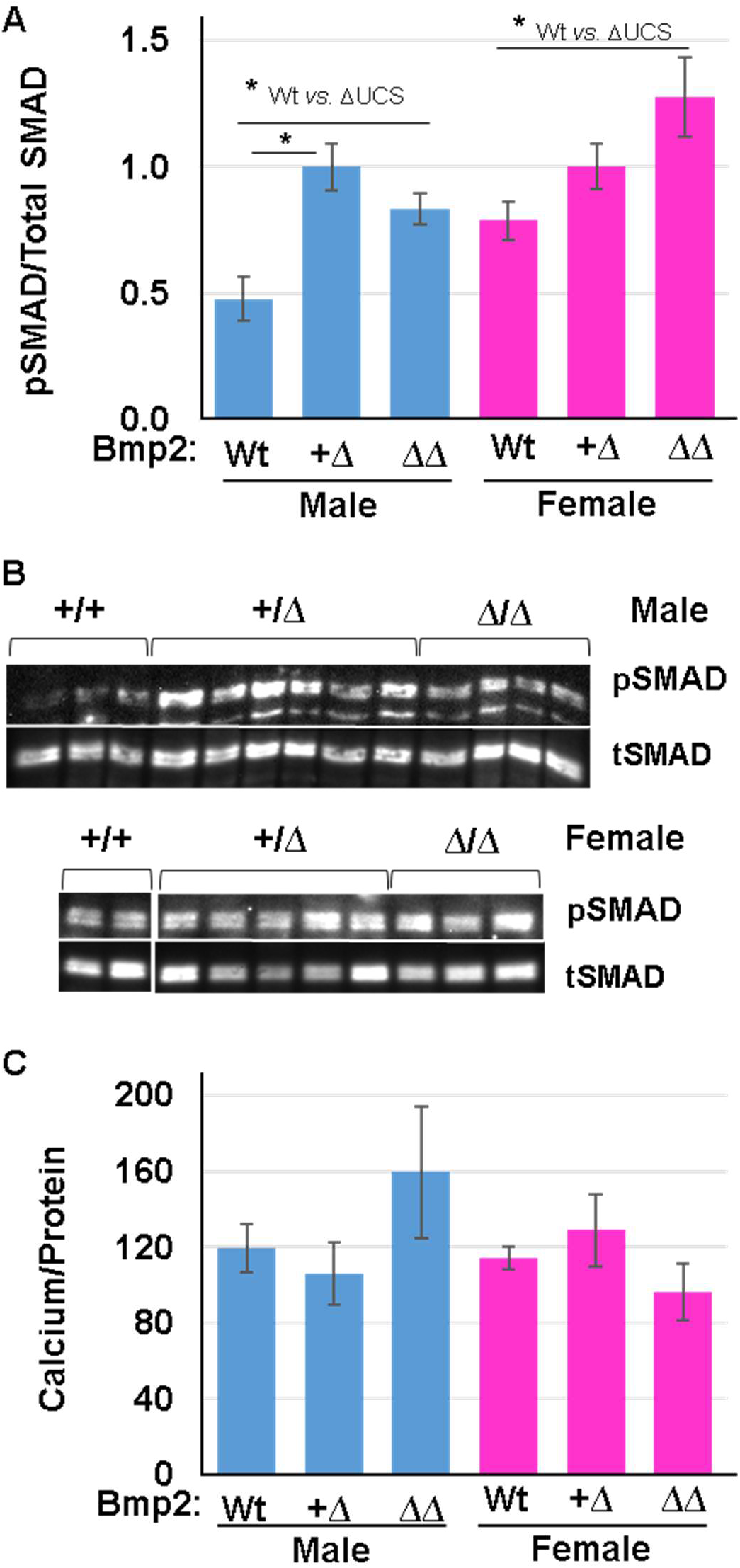
BMP signaling and calcium levels in aorta from *Klotho* mutant mice with *Bmp2* mutations. Aortas were isolated from mice bearing the wild type (Wt), heterozygous (+/Δ), or homozygous UCS deletion (ΔΔ) *Bmp2* genotypes between 40 and 50 days of age. All mice were homozygous for the *Klotho* mutation with renal disease (*Kl*^*kl*/*kl*^). Average parameter values are presented as averages with SEM. **A.** BMP signaling levels as assessed by phosphorylated SMAD1/5/9(8) (pSMAD) levels normalized to total SMAD 1/5/9(8) (tSMAD) (males, n = 3 – 6; females, n = 5 – 6). **B.** Representative western blot panels showing pSMAD1/5/9(8) and total SMAD1/5/9(8) levels, males (top) and females (bottom). **C.** Calcium levels normalized to protein concentration (males, n = 4 – 9; females, n = 5 – 8). All experiments were repeated at least twice with similar results. * *p* < 0.05.

### The *Bmp2 UCS* influences body and heart weights

We observed that the inheritance of UCS deletion allele varied with *Klotho* genotype (Fig. 2D). Therefore, we tested whether deleting the *Bmp2* UCS impacts overall health as assessed by the body weights of the mice. *Bmp2* genotype did not impact the body weight for control animals of both sexes (Fig. 6A). As expected, female control mice weighed about 20% less than males for all 3 *Bmp2* genotypes (*p* = 3.96 × 10^−18^). The severe runting phenotype caused homozygous *Klotho* mutant mice with the wild type *Bmp2* genotype to weigh about a third that of the control mice. Curiously, the typical male to female weight ratio was reversed in *Klotho* mutant homozygotes. Specifically, female *Klotho* homozygotes with the wild type *Bmp2* genotype weighed about 22% more than male homozygotes (Fig. 6A, *p* = 0.04).

**Figure 6.**
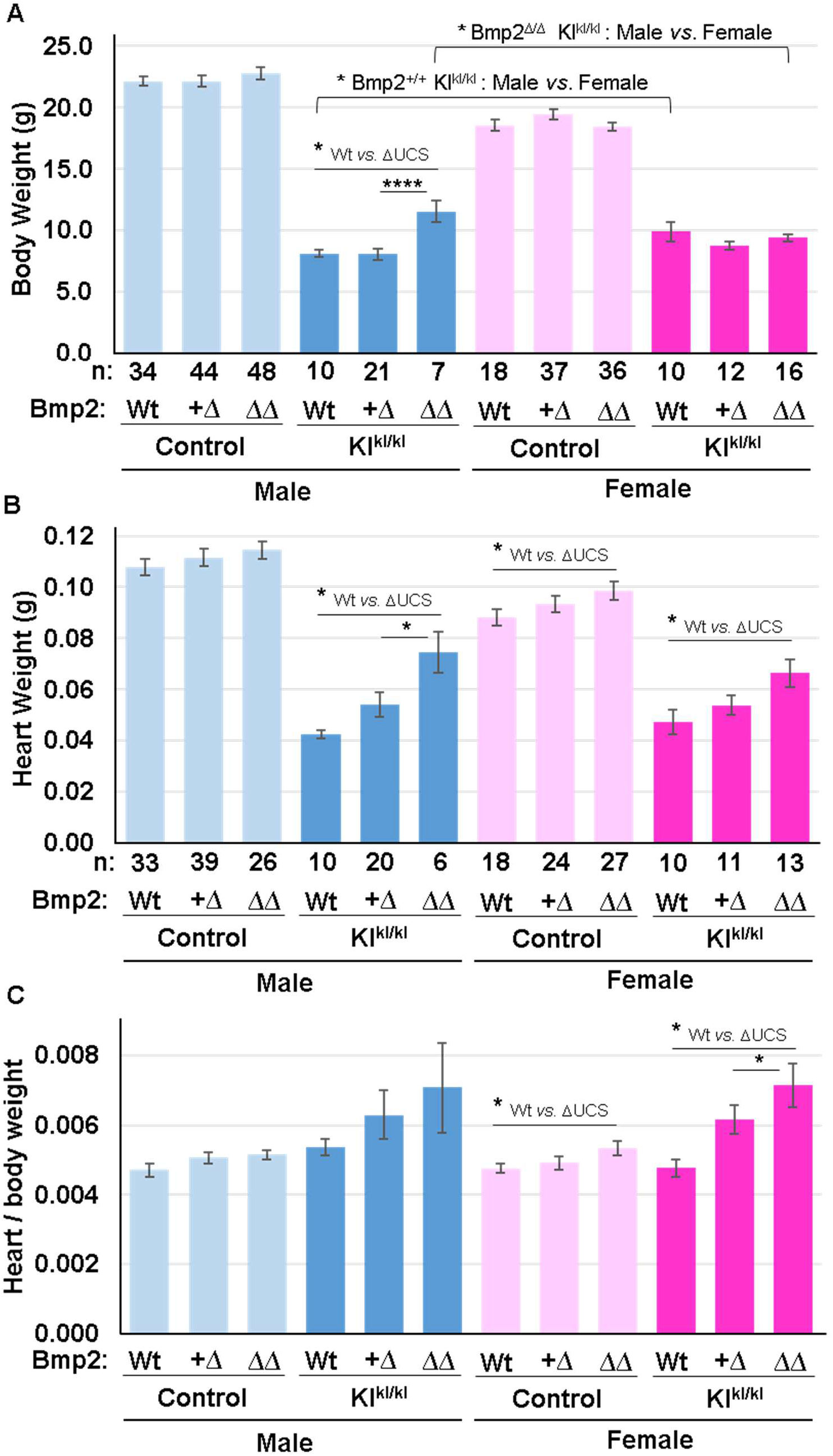
Body and heart weights from healthy control and *Klotho* mutant mice with *Bmp2* mutations. Body **(A)** and heart weights **(B)** from control mice with normal kidney function (*Kl*^+/+^ and *Kl*^*kl*/+^) or diseased mice homozygous for the *Klotho* mutation (*Kl*^*kl*/*kl*^) bearing the wild type (Wt), heterozygous (+/Δ), or homozygous UCS deletion (ΔΔ) *Bmp2* genotypes. Mice were of age 47.1 ± 6 days. Heart weight was normalized to body weight in C. Parameter values are presented as averages with SEM. The number of animals is indicated under the respective bars. * *p* < 0.05, ** *p* < 0.01, \*\*\**p* < 0.005, **** *p* < 0.001.

Homozygous deletion of the UCS (*Bmp2*^Δ*UCS*/Δ*UCS*^) increased the weight of male *Klotho* homozygotes by 40% (Fig. 6A, *p* = 0.0003). Because UCS deletion failed to impact females similarly, the mice lacking the *Bmp2* UCS exhibited the usual male to female weight ratio with males weighing 23% more than females (Fig. 6A, *p* = 0.008). These results suggest a sex-specific difference in the impact of *Bmp2* UCS on the overall physiology of *Klotho* homozygotes. To test if this *Bmp2* allele influenced overall cardiovascular health, we next compared the heart weights of male and female mice with all *Bmp2* and *Klotho* genotypes (Fig. 6B). A modest increase in the heart weights of control mice was associated with UCS deletion. Because heart weight is normally proportional to body weight, we calculated relative heart weights. The small, but significant, increase in the heart weight of female *Klotho* controls lacking the UCS was retained after normalizing to body weight (Fig. 6C, *p* = 0.04).

Deletion of the UCS affected absolute heart weight more dramatically in *Klotho* homozygotes of both sexes. In male and female *Klotho* homozygotes, homozygous UCS deletion (*Bmp2*^Δ*UCS*/Δ*UCS*^) significantly increased absolute heart weight by 75% in males (*p* = 0.0002) and 40% in females (*p* = 0.004, Fig. 6B). Female heart weight normalized to body weight also was increased by nearly 50% (p = 0.005, Fig. 6C). In male *Klotho* homozygotes, a trend towards higher relative weight remained (Fig. 6C) despite the increased overall body weight associated with UCS deletion (Fig. 6A). These results suggest that an intact *Bmp2* UCS protects from an enlarged heart.

## Discussion

Our objective was to test the role of an extraordinarily conserved *Bmp2* regulatory element, the UCS, in the aorta from control adult mice and in mice with genetically induced vascular calcification associated with severely reduced renal function. We used a new *Bmp2* allele without the UCS that increased *Bmp2* RNA and BMP signaling in embryos (31). As we observed in embryos, the UCS can repress BMP synthesis in the adult aorta. However, we also discovered that the UCS functions differently between male and female mice. We will first discuss our findings in males and then compare and contrast these findings to those in females. Our previous study using a *lacZ* reporter gene controlled by the distal *Bmp2* promoter and the *Bmp2* 3’UTR showed that Cre-mediated deletion of the *Bmp2* UCS induced robust gene expression in the aorta, coronary vasculature, and cardiac valves (25). However, this reporter gene lacked the proximal promoter, intronic regulatory elements, and long-range cis-regulatory elements. Thus, the reporter gene only partly recapitulated endogenous *Bmp2* gene expression patterns (19,27). Using our new *Bmp2* allele lacking the UCS (*Bmp2*^ΔUCS^), we showed that the UCS represses *Bmp2* mRNA abundance and BMP signaling in embryos (31). In the present study, we demonstrated for the first time that the UCS also represses *Bmp2* RNA abundance and BMP signaling in control adult male aorta (Fig. 3A - C). This observation, which is consistent with all previous studies using reporter genes *in vitro* and *in vivo* (20,24,25,27), indicates that the UCS can restrain this pro-calcific growth factor in healthy cardiovascular tissues.

In contrast to healthy vascular cells, increased levels of *Bmp2* are tightly associated with pathological calcification of the coronary vasculature (5,6,11,12). To understand why this pro-calcific protein is induced in physiologies such as aging and reduced renal function, we also assessed the capacity of the UCS to restrain BMP signaling in aortic tissue undergoing pathological calcification. In mice bearing the wild type *Bmp2* genotype, but homozygous for the hypomorphic *Klotho* mutation, BMP signaling, and calcium levels were increased in the aorta (Fig. 4B - F). This mirrors the increased levels observed in aortic valves from *Klotho* null mice (29). As in control males, the *Bmp2* UCS repressed BMP signaling in the aorta from male *Klotho* mutant mice (Fig. 5A, B). A trend towards increased aortic calcium levels with *Bmp2* UCS deletion was observed in male *Klotho* mutant mice (Fig. 5C). Thus, in both control and diseased male aorta, the UCS limits BMP signaling. Our observations in males correspond with the simple hypothesis that a functional UCS protects against excessive BMP signaling and maybe against calcification.

Interestingly, results in female mice suggest a more complex story. First, basal BMP signaling was 2-fold higher in aorta from control females relative to males (Fig. 3B). Second, unlike in males, the *Bmp2* UCS failed to repress *Bmp2* mRNA and BMP signaling in control non-calcified aorta from females (Fig. 3A - C). Finally, basal aortic calcium was 36% higher in females relative to males (Fig. 3G). However, as in both healthy control and *Klotho* mutant male mice, the UCS did repress aortic BMP signaling in the female *Klotho* homozygotes. This suggests that *Klotho*-associated renal disease stimulates the braking action of the UCS on aortic BMP signaling in females. Potential sex-related differences in *Bmp2* gene regulatory and BMP signaling mechanisms will be further investigated (33).

The female hormone estrogen may play a role in these differences. Indeed, previous findings in cell culture indicate that estrogen directly induces *Bmp2* transcription (34,37). We confirmed that estrogen stimulates *Bmp2* RNA abundance in the intact aorta (Fig. 3F). The differential impact of deleting the UCS suggests additional dissimilarities in the post-transcriptional processes that regulate BMP2 synthesis in males and females. Sex-related differences in *Bmp2* gene regulation may be clinically relevant as significant disparities in incidence, prognosis, and response to treatments for arterial diseases occur between men and women (38–40).

BMPs were defined by their ability to induce bone from mesenchymal tissues (4). Indeed, all major forms of cardiovascular calcium deposition (aortic valve, medial artery, and atherosclerotic calcification) proceed by mechanisms resembling bone formation (5,9,12,42). However, the increased BMP signaling associated with UCS deletion did not lead clearly to a corresponding increase in aortic calcification. Thus, loss of the UCS-mediated repression of *Bmp2* may not be a sufficient trigger to induced aortic calcification. Indeed, this post-transcriptional repression is only one of the numerous mechanisms that control *Bmp2* gene expression and BMP signaling (19). BMP signaling can activate repressors such as the extracellular antagonist NOGGIN, the intracellular repressor SMAD6, and the miRNAs that repress BMP receptors (43–45). Furthermore, a multitude of positive and negative regulators control the differentiation program leading from bone progenitors to committed osteoprogenitors to differentiating osteoblasts and finally to mature osteoblasts that cause mineralization (46). The redundant feedback mechanisms that dampen BMP signaling along with downstream regulators of osteogenesis would buffer the impact of reduced UCS function on vascular calcification.

Our focus in this study was the regulatory impact of the UCS on *Bmp2* gene expression, BMP signaling, and calcification. However, we observed that the UCS deletion allele modified the *Klotho* phenotype more generally and in a manner also distinguished by sex. First, deletion of the UCS significantly increased the weight of male *Klotho* homozygotes, but not female homozygotes (Fig. 6A). A second phenotype was an increase in heart weight. Although most notable in *Klotho* homozygotes of both sexes, the weights of control female hearts were significantly elevated by deletion of the UCS within only 6 -7 weeks of age (Fig. 6B, C). The UCS deletion allele dramatically altered embryonic morphogenesis and viability, but only in a subset of offspring (31). Because the penetrance of severe embryonic malformations correlated with the level of BMP signaling, we proposed that other BMP pathway regulators compensated for deletion of the repressive UCS. The surviving pups – the subjects of this study – are those with adequate regulation. In the survivors, subtler congenital defects, *e.g.*, of the valves, are possible. Indeed, the edema observed in some embryos is consistent with reduced cardiac function (31). The impact of such developmental anomalies may appear sooner in mice subjected to additional cardiovascular stress, such as the renal failure associated with KLOTHO loss of function. The extraordinary conservation of the UCS is consistent with evolutionary drive to maintain BMP signaling within a narrow developmentally tolerated range. If the remaining redundant regulatory mechanisms are severely inadequate, then catastrophic embryonic malformations may occur. Less severe regulatory changes may reveal themselves only in the contexts of aging or pathological stresses.

## Acknowledgments

We thank Drs. Stella Elkabes and Li Ni for the gift of the aorta from ovariectomized females. We also appreciate the part-time assistance of students Annica Tehim (Rutgers New Jersey Medical School), Amy Song (The College of NJ), Irina Kleiman and Risha Patel (Rutgers School of Graduate Studies).

